# BRN2 and MITF together impact AXL expression in melanoma

**DOI:** 10.1101/2020.07.27.223982

**Authors:** Jacinta L. Simmons, Hannah M. Neuendorf, Glen M. Boyle

**Affiliations:** Cancer Drug Mechanisms Group, QIMR Berghofer Medical Research Institute, Brisbane, Queensland, Australia; School of Biomedical Sciences, Faculty of Health, Queensland University of Technology, Brisbane, Queensland, Australia; School of Biomedical Sciences, Faculty of Medicine, University of Queensland, Brisbane, Queensland, Australia; School of Environment and Science, Griffith University, Brisbane, Queensland, Australia

**Keywords:** POU3F2, therapy resistance, transcription factor

## Abstract

The inverse relationship between transcription factor MITF and receptor tyrosine kinase AXL has received much attention recently. It is thought that melanoma tumors showing AXL^high^/MITF^low^ levels are resistant to therapy. We show here that a population of cells within melanoma tumors with extremely high expression of AXL are negative/low for both MITF and the transcription factor BRN2. Depletion of both transcription factors from cultured melanoma cell lines produced an increase in AXL expression greater than depletion of MITF alone. Further, re-expression of BRN2 led to decreased AXL expression, indicating a role for BRN2 in regulation of AXL levels unrelated to effects on MITF level. As AXL has been recognized as a marker of therapy resistance these cells may represent a population of cells responsible for disease relapse and as potential targets for therapeutic treatment.

## 1. Background

Microphthalmia-associated transcription factor (MITF) is a key melanocytic transcription factor with well documented roles in normal melanocytes and during melanoma progression.^[1]^ Its binding motif is found within 20 kb of over 9000 genes with MITF directly linked to regulation of at least 465 genes from a broad range of processes from cellular senescence to mitosis.^[2]^ MITF is often linked to a phenomenon described as phenotype switching where cells reversibly switch between proliferative (MITF expressing) and invasive (reduced MITF level) states during metastasis.^[3]^ The invasive state characterized by loss of MITF may gain expression of the POU domain transcription factor BRN2 (encoded by *POU3F2*).^[4, 5]^

Despite the requirement for both MITF and BRN2 to be present in a tumor population for efficient melanoma metastasis to occur,^[6]^ expression of MITF and BRN2 is generally accepted to be mutually exclusive at the cellular level and inversely correlated in melanoma tumors, xenografts and 3D cultured cells but not 2D culture.^[4, 5, 7]^ The reciprocal relationship between MITF and BRN2 is maintained by both direct binding of BRN2 to the *MITF* promoter and the indirect suppression of BRN2 by MITF through miR-211.^[5, 8]^

Another noteworthy inverse relationship is that which exists between MITF and the receptor tyrosine kinase AXL.^[9]^ This relationship has drawn much interest with AXL^high^/MITF^low^ cells being described as resistant to targeted MAPK inhibitor therapy.^[10, 11]^ It was also reported that invasion in MITF^low^ cells is dependent upon AXL and the transcription factor FRA-1.^[9, 11]^

With MITF having an inverse relationship with both BRN2 and AXL we sought to investigate the relationship between the three proteins further to characterize their contribution to melanoma heterogeneity.

## 2. Questions Addressed

What is the relationship between MITF, AXL and BRN2 in melanoma cells?

## 3. Experimental Design

### 3.1. Analysis of publically available data

The Gerber dataset; GSE81383^[12]^ was downloaded from NCBI’s Gene Expression Omnibus. The normalized read count was converted to log_10_, values of 0 were assigned −2 log_10_ and GraphPad Prism version 8.2.1 used for linear regression analysis. For pooling of data the cut off value between low and high expression was set to 2 for *AXL*, 0.1 for *POU3F2* and 1.6 for *MITF*.

### 3.2. Cell culture

Melanoma cell lines were maintained in RPMI-1640 with 10% FBS (Life Technologies, Carlsbad, CA). Routine mycoplasma tests were always negative and short tandem repeat profiling confirmed cell line identity.

### 3.3. Inducible knockdown and expression

For both knockdown and overexpression cell lines we used the BLOCK-iT Lentiviral expression system (Life Technologies). We have previously described the overexpression of BRN2 and depletion of MITF and BRN2.^[6, 13]^ We have further modified the same system to simultaneously express shRNA’s targeting MITF and BRN2.

### 3.4. Western blot analysis

Cell lysates (30 μg protein) were resolved by SDS-PAGE and transferred to nitrocellulose membranes. Antibodies used in this study were: anti-MITF (12590; Cell Signaling Technologies [CST], Danvers, MA), anti-BRN2 antibody (12137; CST), anti-AXL antibody (8661; CST), anti-GAPDH (R&D Systems, Minneapolis, MN). Where required, bands were quantified by densitometry using ImageJ version 1.52a and analyzed by linear regression analysis using GraphPad Prism version 8.2.1 (GraphPad Software, San Diego, CA).

### 3.5. Quantitative reverse transcription PCR

RNA was extracted from cells using the QIAGEN RNeasy plus kit (Hilden, Germany) and 2 μg of total RNA was reverse transcribed with SuperScript III reverse transcriptase (Invitrogen, Carlsbad, CA) following the manufacturer’s protocol. The CFX384 Real-Time System (BioRad Laboratories, Hercules, CA) was used for qRT-PCR with SYBR Green PCR master mix (Applied Biosystems, Foster City, CA). The *AXL* primer sequences were (5’ - 3’); Forward: ACCTACTCTGGCTCCAGGATG; Reverse: CGCAGGAGAAAGAGGATGTC. *GAPDH* primers have been described previously.^[6]^

## 4. Results

To investigate the relationship between MITF, BRN2 and AXL, we used a publically available gene expression dataset to explore expression correlations between *MITF, BRN2 (POU3F2)* and *AXL*. The Gerber dataset (GSE81383) consists of single cell RNA-seq of 307 cells from three early passage melanoma patient biopsies.^[12]^ An inverse correlation between *MITF* and *AXL* in this dataset has been reported previously.^[14]^ Similarly, we found a strong inverse relationship between *MITF* and *AXL* (Figure 1a, R^2^ = 0.5648, *P* = <0.0001) and between *MITF* and *POU3F2* (Figure 1b, R^2^ = 0.2127, *P* = <0.0001) using linear regression analysis of the log_10_ normalized read count (read count values of 0 were arbitrarily assigned a value of −2). Interestingly a positive correlation was also seen between *AXL* and *POU3F2* (Figure 1c, R^2^ = 0.3562, *P* = <0.0001).

**Figure 1.**
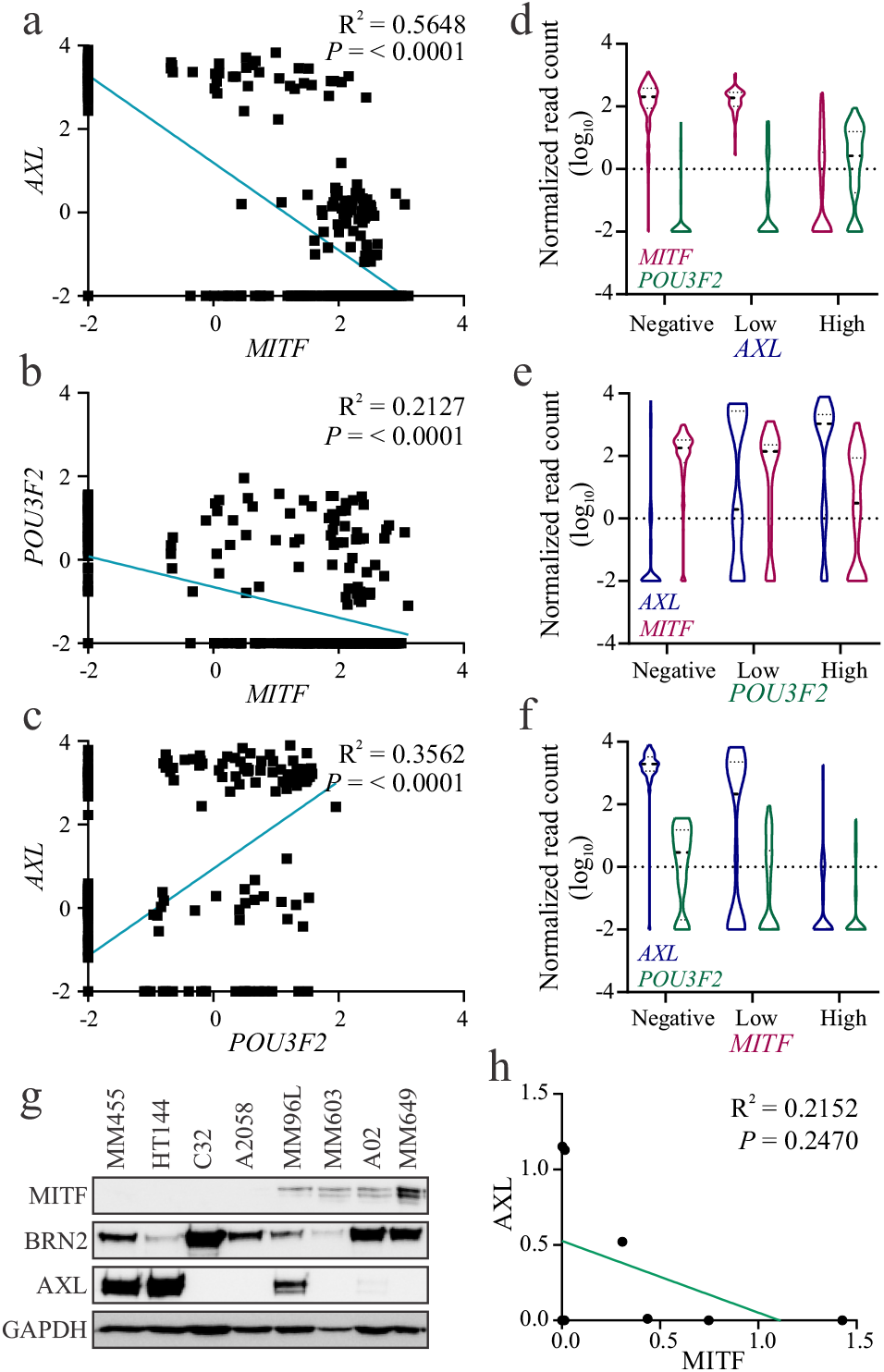
Heterogeneous expression of *MITF*, *POU3F2* and *AXL* in single melanoma cells (a-c) Linear regression analysis of *MITF, POU3F2 and AXL* expression in the Gerber dataset (GSE81383) from single cell RNA sequencing of short term patient biopsy cultures.^[12]^ Normalized read count is expressed as log_10_ and values of 0 were assigned as 0.01 (−2 log_10_). (d-f) Separation of cells on the basis of the log_10_ normalized read count into negative, low and high groups. Negative = −2; low = < 2 (d, *AXL*, blue), < 0.1 (e, *POU3F2*, green) or < 1.6 (f, *MITF*, pink); high = > 2 (d, *AXL*), > 0.1 (e, *POU3F2*) or > 1.6 (f, *MITF*). Broken line = median, dotted lines = quartiles (g) Western blot showing expression of MITF, BRN2 and AXL in cultured melanoma cell lines. (h) Quantification of western blot shown in panel g, and analysis of correlation between MITF and AXL by linear regression analysis (R^2^ = 0.2152, *P* = 0.2470).

The data showed that cells appeared to cluster into three distinct groups on the basis of AXL expression (Figure 1a, c). We therefore grouped data on the basis of AXL expression (negative, low, high) and graphed the log_10_ normalized read count of *MITF* and *POU3F2* (Figure 1d). Interestingly, while *AXL* negative or low cells maintained the inverse relationship between *MITF* and *POU3F2* this was lost in *AXL* high cells. Similarly, separation of cells into three groups on the basis of *POU3F2* expression showed an inverse relationship between *MITF* and *AXL* in the *POU3F2* negative group only (Figure 1e). These results highlight the complexity of the relationship between these three proteins.

Importantly, *MITF* and *POU3F2* expression is not entirely mutually exclusive, even at single cell level.^[15]^ A proportion of *MITF* negative cells were also negative for *POU3F2* while cells expressing high levels of *POU3F2* had great variability in the level of *MITF* expression (Figure 1e, f). We also noted a population of cells that had high expression of *AXL* and low or negative expression of both *MITF* and *POU3F2* (Figure 1d). This was further evident when *AXL* high cells were graphed with connecting lines and colored on the basis of *POU3F2* expression (Figure S1).

Western blot analysis of a panel of 8 metastatic melanoma derived cell lines (Figure 1g) revealed an inverse relationship between MITF and AXL (Figure 1h) although due to low sample numbers this was not statistically significant. No correlation was seen between MITF and BRN2 or AXL and BRN2 (Figure S2a, b). However, the limitation of the western blot analysis is that levels of MITF and BRN2 are known to vary between cells grown in 2D culture and 3D cultured cells or patient samples.^[16]^

Given the results from single cell RNAseq where a proportion of AXL high cells were negative for MITF and BRN2 expression, we wished to examine the effect of reduction of these transcription factors on AXL expression. Using doxycycline inducible shRNA expression we were able to deplete MITF and BRN2 in two metastatic melanoma cell lines (A02 and MM649) by the addition of 50 ng/mL doxycycline to culture media for 96 h (Figure 2a, b). AXL protein level increased upon depletion of MITF either alone or combined with depletion of BRN2. Interestingly, the level of AXL was much higher in cells depleted of both BRN2 and MITF when compared to cells depleted of MITF alone (Figure 2a, b). Analysis of the same cell lines by qRT-PCR showed a significant increase in *AXL* transcript when MITF and BRN2 were depleted in tandem compared to only MITF depletion (Figure 2c, *P* = 0.0393, d, *P* = 0.0109). In addition, MM96Lc8LC cells, a line negative for constitutive expression of both BRN2 and MITF^[16, 17]^ also showed an upregulation of AXL protein level compared to the control cells that express both transcription factors (MM96Lc8D) (Figure 2e). Further, re-expression of BRN2 in metastatic melanoma cell lines resulted in decreased AXL expression but variable changes to MITF expression (Figure 2f, g). Both cell lines have extremely low expression of MITF (allowing expression of AXL) however in MM455 cells MITF expression increased slightly on expression of BRN2 (Figure 2f) while the reverse was true for D04 (Figure 2g).

**Figure 2.**
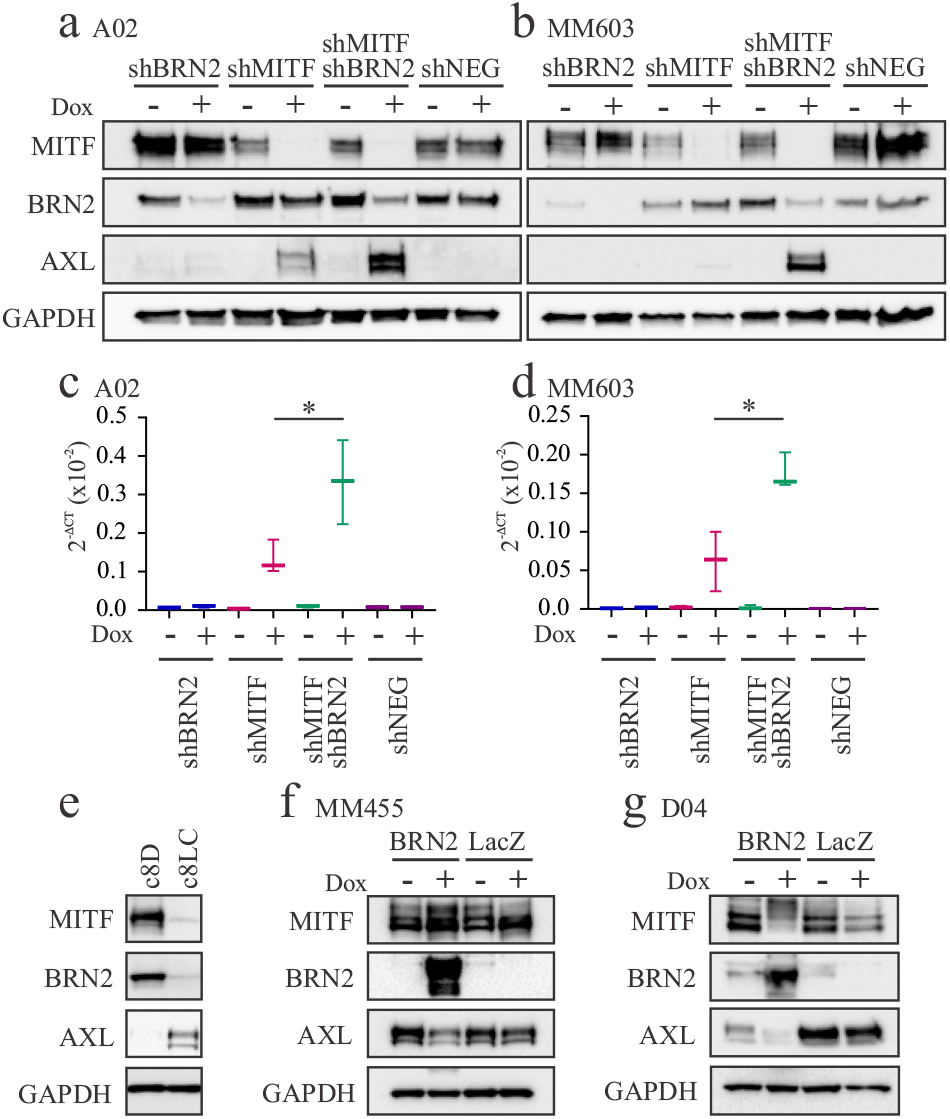
BRN2, as well as MITF, impacts AXL expression. Western blot analysis of MITF, BRN2 and AXL expression in A02 (a) and MM603 (b) melanoma cell lines expressing shRNA against MITF, BRN2 or both. Analysis of *AXL* expression by qRT-PCR in (c) A02 and (d) MM603 cells expressing shRNA as indicated, data represents mean and SD of 3 biological replicates. \**P* < 0.05, paired t-test (e) Western blot analysis of MITF, BRN2 and AXL in clones of the MM96L cell line; MM96Lc8LC (c8LC) and MM96Lc8D (c8D). Western blot analysis of inducible BRN2 expression in (f) MM455 and (g) D04 melanoma cell lines.

Taken together these results indicate a role for both MITF and BRN2 in the transcriptional regulation of *AXL*. The relationship between MITF and AXL in melanoma is well reported.^[9–11]^ Interestingly, expression of MITF in immortalized melanocytes caused increased expression of AXL.^[18]^ Mechanistically, it appears that neither MITF nor BRN2 directly repress AXL expression. Analysis of the promoter region of AXL as well as published ChIP-seq data do not show potential binding of either transcription factor. However, indirect control of AXL level may be mediated by induction of miRNA expression by MITF and BRN2. MITF has been reported to induce expression of miR-363^[19]^ while BRN2 binds at the promoter regions of miR-101 and miR-103^[20]^; binding sites for each miRNA are present in the *AXL* 3’ UTR. Further experimental validation of these mechanistic relationships is required.

## 5. Conclusions

These results highlight that in the extremely heterogeneous melanoma tumor environment protein expression patterns are tremendously complicated. Traditional ‘bulk’ analysis of expression within a tumor could result in only the major contributing cell populations being identified. This was highlighted in a recent study where two different MITF^low^ populations were identified in a patient-derived xenograft model of minimal residual disease.^[21]^ One population was associated with an EMT signature and was decreased in residual disease (compared to untreated) while the de-differentiation associated population was enriched at the same stage. Crucially, as upregulation of AXL is currently recognized as a marker of resistance to therapy the presence of MITF^low^/BRN2^low^/AXL^high^ cells could be responsible for disease relapse, or indicate a requirement for closer monitoring of patients during treatment.

## DATA AVAILABILITY

All data used in the current study are available from the Gene Expression Omnibus (GEO), number GSE81383. Accessible from https://www.ncbi.nlm.nih.gov/geo/

## ACKNOWLEDGEMENTS

The authors wish to thank the National Health and Medical Research Council of Australia (APP1158283 Project Grant) for support.

## AUTHOR CONTRIBUTIONS

JLS and GMB designed the study. JLS and HMN performed the research. JLS and GMB analyzed the data and wrote the manuscript. All authors revised the manuscript.

## REFERENCES

[1] C. R. Goding and H. Arnheiter, Genes Dev 2019, 33, 983–1007.

[2] T. Strub, S. Giuliano, T. Ye, C. Bonet, C. Keime, D. Kobi, S. Le Gras, M. Cormont, R. Ballotti, C. Bertolotto and I. Davidson, Oncogene 2011, 30, 2319–2332.

[3] K. S. Hoek, O. M. Eichhoff, N. C. Schlegel, U. Döbbeling, N. Kobert, L. Schaerer, S. Hemmi and R. Dummer, Cancer Res 2008, 68, 650–656.

[4] S. Pinner, P. Jordan, K. Sharrock, L. Bazley, L. Collinson, R. Marais, E. Bonvin, C. Goding and E. Sahai, Cancer Research 2009, 69, 7969–7977.

[5] J. Goodall, S. Carreira, L. Denat, D. Kobi, I. Davidson, P. Nuciforo, R. A. Sturm, L. Larue and C. R. Goding, Cancer Research 2008, 68, 7788–7794.

[6] J. L. Simmons, C. J. Pierce, F. Al-Ejeh and G. M. Boyle, Sci Rep 2017, 7, 10909.

[7] M. Ennen, C. Keime, D. Kobi, G. Mengus, D. Lipsker, C. Thibault-Carpentier and I. Davidson, Oncogene 2015, 34, 3251–3263.

[8] G. M. Boyle, S. L. Woods, V. F. Bonazzi, M. S. Stark, E. Hacker, L. G. Aoude, K. Dutton-Regester, A. L. Cook, R. A. Sturm and N. K. Hayward, Pigment Cell Melanoma Res 2011, 24, 525–537.

[9] M. Sensi, M. Catani, G. Castellano, G. Nicolini, F. Alciato, G. Tragni, G. De Santis, I. Bersani, G. Avanzi, A. Tomassetti, S. Canevari and A. Anichini, J Invest Dermatol 2011, 131, 2448–2457.

[10] D. J. Konieczkowski, C. M. Johannessen, O. Abudayyeh, J. W. Kim, Z. A. Cooper, A. Piris, D. T. Frederick, M. Barzily-Rokni, R. Straussman, R. Haq, D. E. Fisher, J. P. Mesirov, W. C. Hahn, K. T. Flaherty, J. A. Wargo, P. Tamayo and L. A. Garraway, Cancer Discov 2014, 4, 816–827.

[11] J. Muller, O. Krijgsman, J. Tsoi, L. Robert, W. Hugo, C. Song, X. Kong, P. A. Possik, P. D. Cornelissen-Steijger, M. H. Foppen, K. Kemper, C. R. Goding, U. McDermott, C. Blank, J. Haanen, T. G. Graeber, A. Ribas, R. S. Lo and D. S. Peeper, Nat Commun 2014, 5, 5712.

[12] T. Gerber, E. Willscher, H. Loeffler-Wirth, L. Hopp, D. Schadendorf, M. Schartl, U. Anderegg, G. Camp, B. Treutlein, H. Binder and M. Kunz, Oncotarget 2017, 8, 846–862.

[13] C. J. Pierce, J. L. Simmons, N. Broit, D. Karunarathne, M. F. Ng and G. M. Boyle, Oncogenesis 2020, 9, 64.

[14] H. Loeffler-Wirth, H. Binder, E. Willscher, T. Gerber and M. Kunz, Biology (Basel) 2018, 7.

[15] M. Ennen, C. Keime, G. Gambi, A. Kieny, S. Coassolo, C. Thibault-Carpentier, F. Margerin-Schaller, G. Davidson, C. Vagne, D. Lipsker and I. Davidson, Clin Cancer Res 2017, 23, 7097–7107.

[16] A. E. Thurber, G. Douglas, E. C. Sturm, S. E. Zabierowski, D. J. Smit, S. N. Ramakrishnan, E. Hacker, J. H. Leonard, M. Herlyn and R. A. Sturm, Oncogene 2011, 30, 3036–3048.

[17] J. A. Thomson, K. Murphy, E. Baker, G. R. Sutherland, P. G. Parsons and R. A. Sturm, Oncogene 1995, 11, 691–700.

[18] T. J. Lavelle, T. N. Alver, K. M. Heintz, P. Wernhoff, V. Nygaard, S. Nakken, G. F. Øy, L. Bøe, A. Urbanucci and E. Hovig, Cancers (Basel) 2020, 12.

[19] F. Ozsolak, L. L. Poling, Z. Wang, H. Liu, X. S. Liu, R. G. Roeder, X. Zhang, J. S. Song and D. E. Fisher, Genes Dev 2008, 22, 3172–3183.

[20] D. Kobi, A. L. Steunou, D. Dembele, S. Legras, L. Larue, L. Nieto and I. Davidson, Pigment Cell Melanoma Res 2010, 23, 404–418.

[21] F. Rambow, A. Rogiers, O. Marin-Bejar, S. Aibar, J. Femel, M. Dewaele, P. Karras, D. Brown, Y. H. Chang, M. Debiec-Rychter, C. Adriaens, E. Radaelli, P. Wolter, O. Bechter, R. Dummer, M. Levesque, A. Piris, D. T. Frederick, G. Boland, K. T. Flaherty, J. van den Oord, Voet, S. Aerts, A. W. Lund and J. C. Marine, Cell 2018, 174, 843–855.e819.

