## Supplementary Figures for "BRN2 and MITF together impact AXL expression in melanoma"

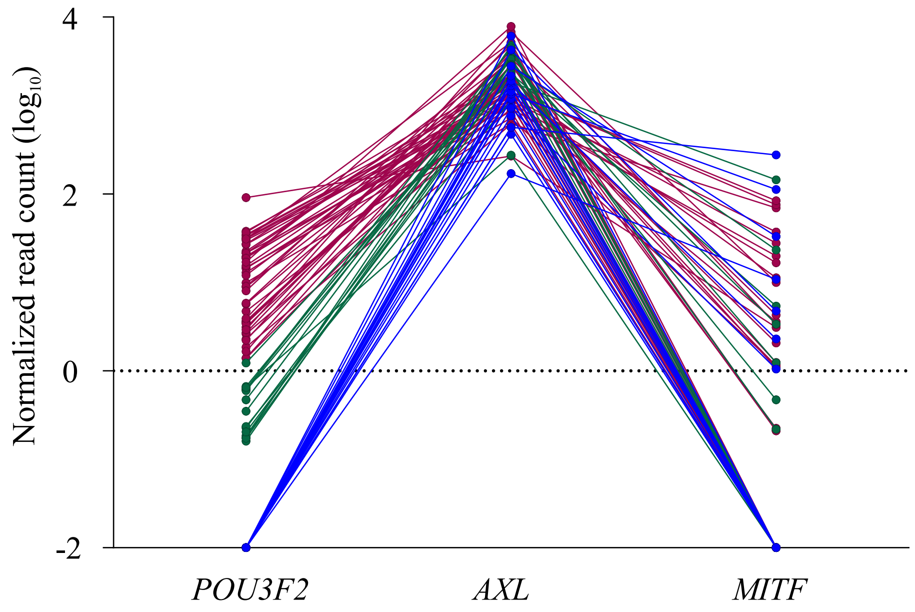


**Figure S1.** Melanoma tumors contain a population of *AXL* high cells with low/no expression of *MITF* and *POU3F2*. The log10 normalized read count for *AXL* high cells from the Gerber dataset was graphed. Single cells are connected by lines. Colors represent *POU3F2* expression, blue = negative, green = low, pink = high.

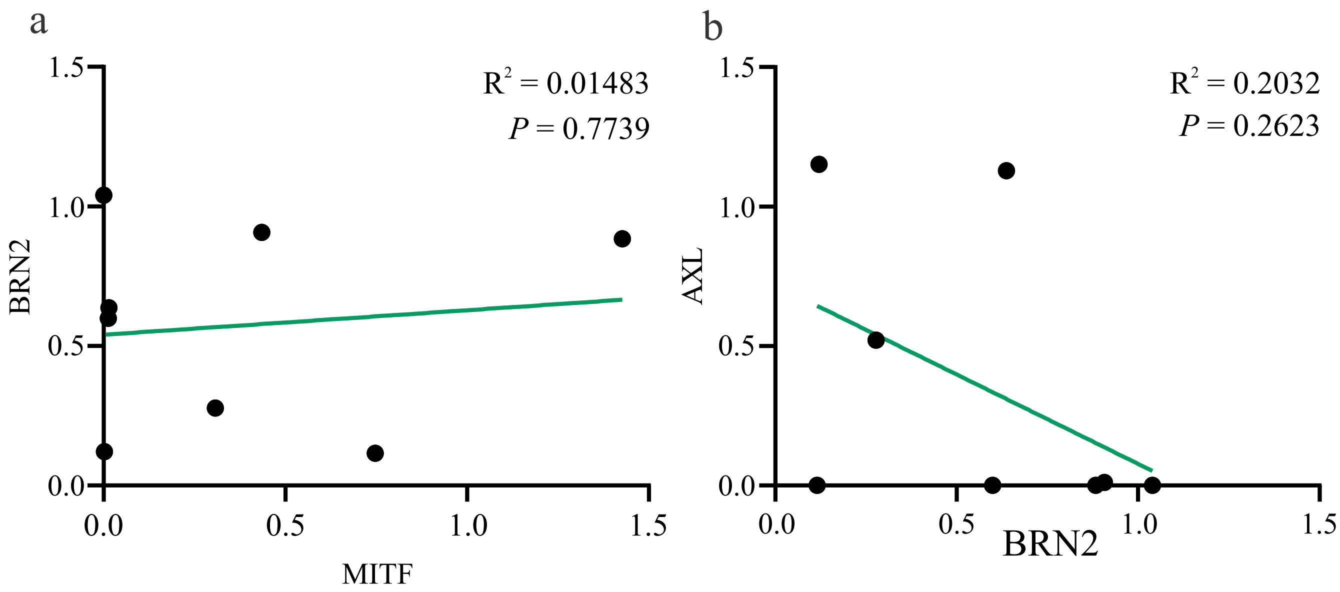


**Figure S2.** Western blots of MITF, BRN2 and AXL expression in melanoma cell lines were quantified by densitometry and analyzed by linear regression analysis. No correlation was found between (a) MITF and BRN2 ($R^2 = 0.01483$, $P = 0.7739$) or (b) AXL and BRN2 ($R^2 = 0.2032$, $P = 0.2623$).
